# plot2DO: a tool to assess the quality and distribution of genomic data

**DOI:** 10.1101/189449

**Authors:** Răzvan V. Chereji

## Abstract

**Summary:** Micrococcal nuclease digestion followed by deep sequencing (MNase-seq) is the most used method to investigate nucleosome organization on a genome-wide scale. We present *plot2DO*, a software package for creating 2D occupancy plots, which allows biologists to evaluate the quality of MNase-seq data and to visualize the distribution of nucleosomes near the functional regions of the genome (*e.g*. gene promoters, origins of replication, etc.).

**Availability And Implementation:** The *plot2DO* open source package is freely available on GitHub at https://github.com/rchereji/plot2DO under the MIT license.

**Supplementary Information:** Supplementary data are available at *Bioinformatics* online.

## 1 Introduction

Nucleosomes - 147 bp of DNA wrapped around a histone octamer in about 1.7 turns - are the basic units of DNA packaging in eukaryotes (Luger, et al., 1997). Access of DNA-binding factors to their target sites is about 10-20 times higher if these sites are located in nucleosome free regions (NFRs) (Liu, et al., 2006). Therefore, knowing the precise positions of nucleosomes and NFRs is very important for understanding DNA-binding and gene regulation.

Currently, the most used method for mapping nucleosomes is MNase-seq: chromatin is digested with micrococcal nuclease (MNase) and the remaining undigested DNA fragments are subjected to high-throughput sequencing. Unfortunately, MNase has a strong sequence preference (Dingwall, et al., 1981; Hörz and Altenburger, 1981), and the nucleosomal fragments that result from an MNase-seq experiment are affected by the degree of MNase digestion (Chereji, et al., 2016; Chereji, et al., 2017). Furthermore, after a mild digestion, a large fraction of the genome is not yet broken into mono-nucleosomal DNA fragments (~150bp long) and is discarded from further analysis, while after an extensive digestion, many nucleosomes occupying A/T-rich sequences are over-digested and lost from the sample of mono-nucleosomal fragments (Chereji, et al., 2017). Therefore, MNase-seq experiments require a careful control of the level of digestion, and the variable degree of digestion must always be taken into account, especially when multiple samples are compared, and differences in nucleosome occupancy are observed among the samples. Here, we present *plot2DO*, a tool for plotting the 2D occupancy (2DO) of genomic data, which is extremely useful not only to assess the degree of digestion as an initial quality check of MNase-seq data, but also for getting insights about the nucleosome organization near functional regions of the genome and about the MNase digestion kinetics.

## 2 Usage

*Plot2DO* is an open source package written in R, which can be launched from the command line in a terminal. The user selects the type of distribution to be plotted (occupancy/coverage of undigested DNA fragments, distribution of fragment centers (nucleosome dyads), or distribution of the 5’/3’ ends of the fragments), the reference points to be aligned (transcription start sites (TSS), transcript termination sites (TTS), +1 nucleosomes, or a list of specific user-provided sites). The user can choose the width of the window that is plotted and can also perform *in silico* size selection of the undigested DNA, by specifying the size limits of the fragments to be used as representative for the nucleosome population.

*Plot2DO* allows the investigation of paired-end sequencing data originating from a variety of organisms (yeast, fly, worm, mouse, and human) and mapped to any of the following genome versions: sacCer3, dm3, dm6, ce10, ce11, mm9, mm10, hg18, hg19. The full list of available options and multiple usage examples are provided as supplementary data at *Bioinformatics* online.

## 3 Discussion and conclusion

The usefulness of *plot2DO* is demonstrated in the figure above. Fig. 1A shows the default panels generated by *plot2DO*. The 2DO plot (heat map) indicates the relative coverage of the DNA fragments of specified lengths, at different locations relative to the selected reference (TSS by default). Each row of the 2DO plot shows the average occupancy generated by DNA fragments of a given length (indicated on the right side) as a function of the position (indicated on top). Each column of the 2DO plot shows the average occupancy generated at a specific position relative to the reference point, as a function of the DNA fragment length. The average one-dimensional occupancy (shown above the 2DO plot), generated by stacking DNA fragments of all lengths, represents the sum of the elements in each column of the matrix shown in the heat map. To compute these occupancy profiles, the raw sequencing data is normalized such that the average occupancy is 1, for each chromosome. The third panel that is generated by *plot2DO* is the fragment length histogram, which is shown to the right of the 2DO plot. Note that this histogram takes into account all the sequencing reads, not just the ones that are mapped to the regions shown in the heat map (*i.e*. the lengths of the reads far from the reference points are also considered).

**Fig. 1.**
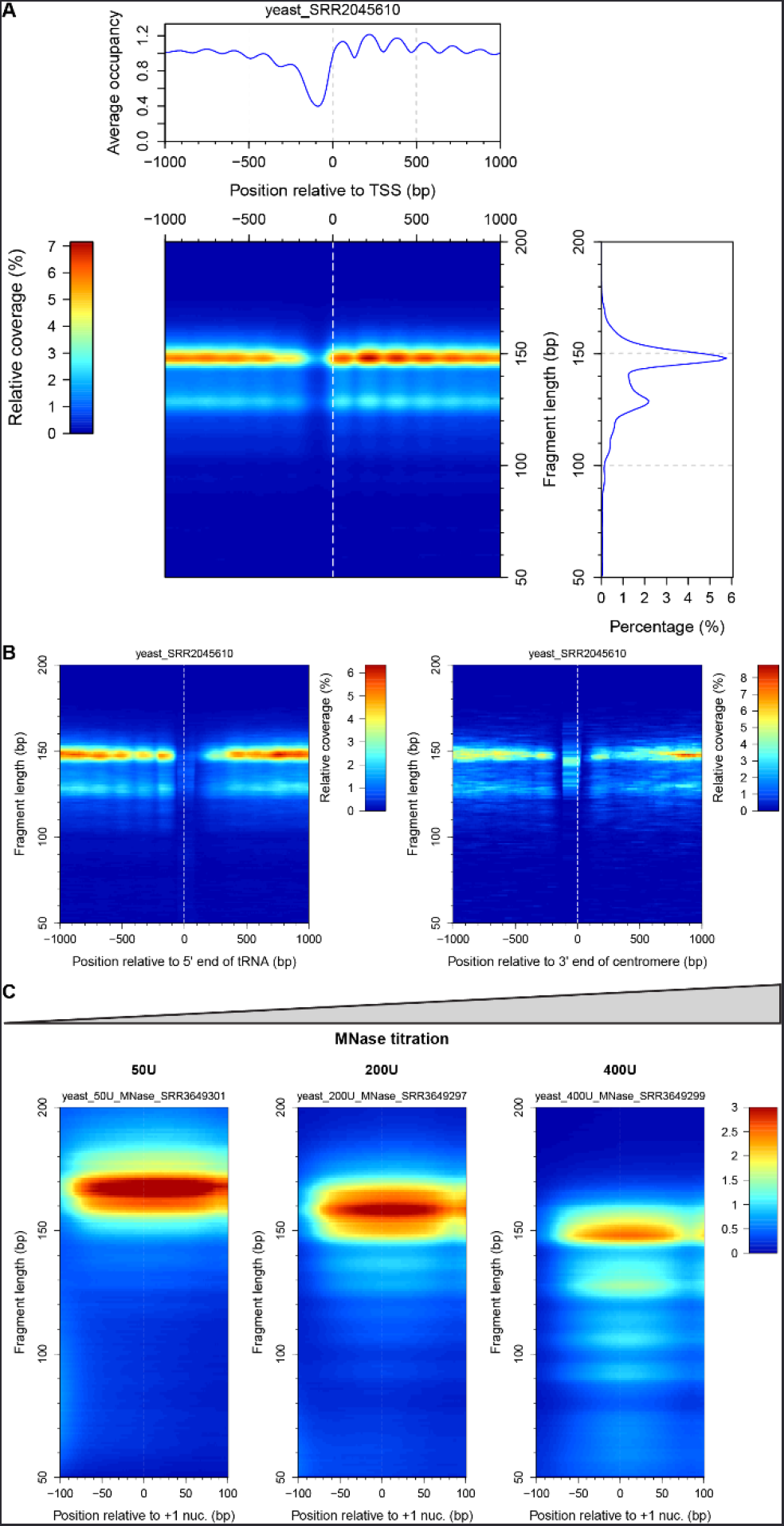
(A) Default figure generated by plot2DO showing the distribution of sequencing reads near the yeast TSSs. (B) *Plot2DO* can also align specific lists of sites, provided by the user, as tRNA genes (left) and centromeric regions (right). (C) Titration of MNase illustrating the effect of different degrees of digestion to the sample of +1 nucleosomes that are recovered in the experiment. For details and usage examples, see the Supplementary information.

Apart from the TSS/TTS alignments, *plot2DO* can also align specific lists of sites that are provided by the user (Fig. 1B). The plots created by *plot2DO* are particularly useful for investigating the level of digestion and the effect of MNase on the subset of intact nucleosomes that are obtained in a sample. Fig. 1C shows the effect of MNase on the +1 nucleosomes (the first nucleosomes downstream from the gene promoters). It is obvious that during the MNase digestion, the fragments that are protected by nucleosomes become shorter and are eventually destroyed by MNase, if the digestion is not stopped early enough. Data used in Fig. 1 were obtained from (Ocampo, et al., 2016) and (Chereji, et al., 2017) (see the Supplementary information for details).

Nucleosome mapping experiments are very expensive, and it is very important to assess the quality of MNase digestion before a large amount of money is spent for sequencing. We recommend that before committing to a large sequencing experiment that results in hundreds of millions of reads, a trial sequencing round should be performed (*e.g*. getting only a few million reads) and the level of digestion should be examined using *plot2DO*. We strongly advocate that in all MNase-seq and MNase-ChIP-seq experiments, *plot2DO* should be the first *plot to do*.

## Acknowledgements

We thank Natalia Petrenko, Ming-an Sun, and Yashpal Rawal for insightful discussions and for testing the software.

## Funding

This work has been supported by the Intramural Research Program of the National Institutes of Health (NIH), and utilized the computational resources of the NIH HPC Biowulf cluster (http://hpc.nih.gov).

## Conflict of Interest

none declared.

## Supplementary Information

*plot,2DO:* a tool to clSSGSS the quality and distribution of genomic data Razvan V. Chereji Division of Developmental Biology, *Eunice Kennedy Shriver* National Institute for Child Health and Human Development, National Institutes of Health, Bethesda,

Maryland 20892, USA

### Download

The easiest way to download this package is by using the GitHub interface (click the green “Clone or download” button) from https://github.com/rchereji/plot2D0.

If you want to download the package from the terminal, then you need to install first a git client of your choice, using the package manager that is available for your system. For example, in Ubuntu or other Debian-based distributions of Linux, you can use apt-get:

~~~
$ sudo apt—get install git
~~~

In Fedora, CentOS, or Red Hat Linux, you can use `yum`:

~~~
$ sudo yum install git
~~~

Installers for Windows and OSX are available for download at the following websites: http://git-scm.com/download/win and http://git-scm.com/download/mac.

After git has been installed, run the following command from the folder where you want to download the *plot2DO* package:

~~~
$ git clone https://github.com/rchereji/plot2DO.git
~~~

In order to be able to run *plot2DO*, you will need to have R installed on your computer, together with the following R packages.

### Dependencies

*Plot2DO* uses the following R packages: biomaRt, caTools, colorRamps, GenomicRanges, optparse, rtracklayer, and Rsamtools. To install these packages, open R and execute the following commands:

~~~
# Install caTools, colorRamps and optparse packages from CRAN:
inst all . packages (c (“caTools” , “colorRamps”, “optparse“))
# Get the latest version of Bioconductor:
source(“https://bioconductor.org/biocLite.R“)
biocLite ()
# Install the remaining Bioconductor packages:
biocLite (c (“biomaRt “ , “GenomicRanges”, “rtracklayer”, “ Rsamtools”))
~~~

### Usage examples

After the R packages have been installed, you can execute *plot2DO* from a terminal. The only required input is the name of the file that contains the aligned genomic data (in BAM or BED format). To generate the default 2D occupancy (2DO) plot (around TSS, all DNA fragments with the lengths between 50 - 200 bp, 2kb windows centered on TSS, yeast DNA), run one of following commands:

~~~
$ Rscript plot2DO.R —fi1e=yeast_SRR2045610.bam               #long version of the flag (—fi1e =<FILENAME>)
$ Rscript plot2DO.R -f yeast_SRR2045610.bam                         # short version of the flag (-f <FILENAME>)
~~~

where `yeast_SRR2045610.bam` is a test BAM file containing paired-end reads, which were aligned to the yeast genome. This command will generate Figure S1.

**Supplementary Figure S1.**
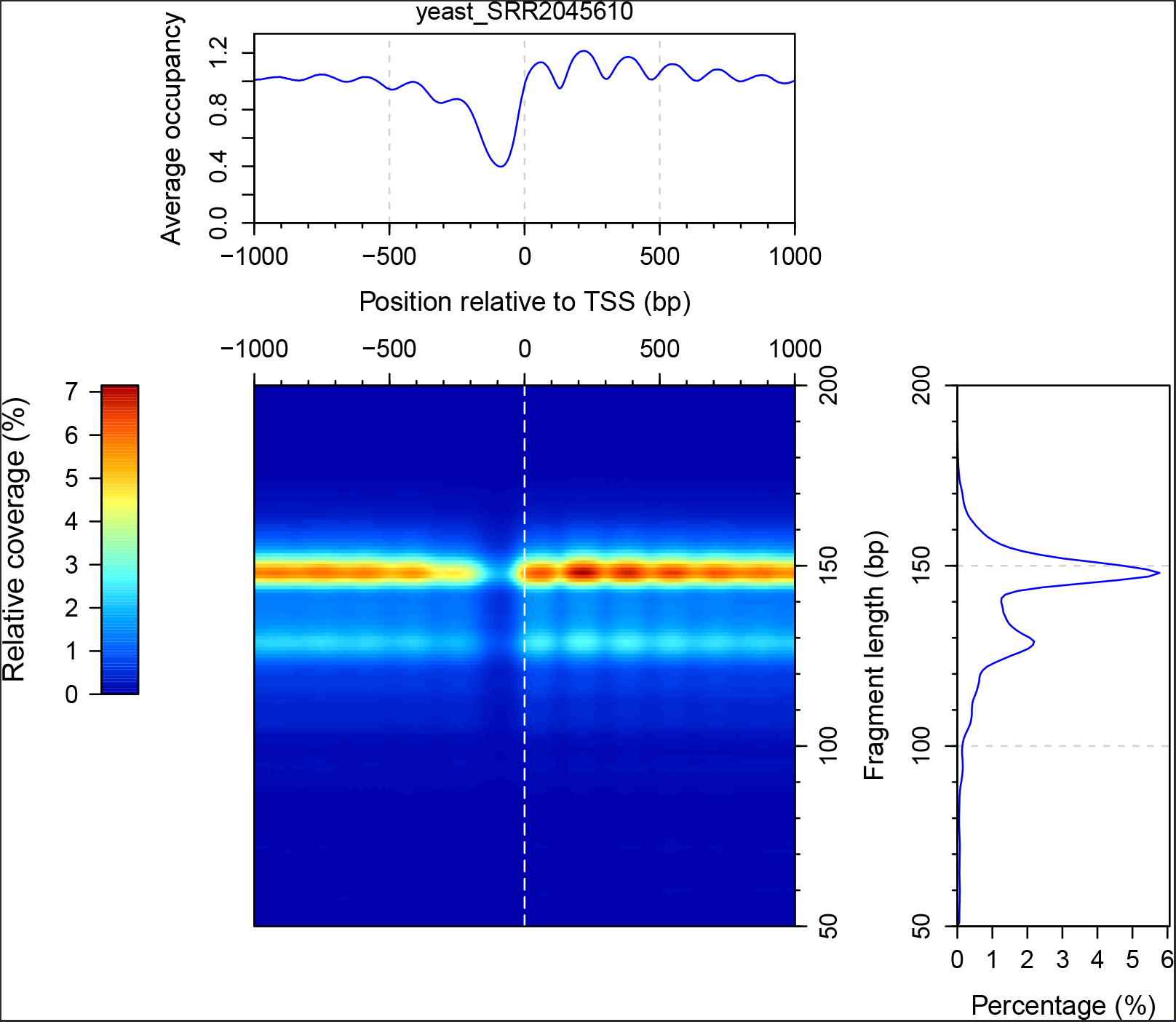
The three panels that are generated by *plot2DO*. (1) The 2D occupancy (2DO) plot -heat map indicating the relative coverage of the DNA fragments of specified lengths, at different locations relative to the reference/alignment points (transcription start sites, or TSSs, in this case). The color red indicates a high coverage, while dark blue indicates zero coverage. (2) The one-dimensional occupancy/coverage (top panel), generated by stacking DNA fragments of all lengths shown in the heat map. (3) Fragment length histogram (right panel) - indicating the percentage corresponding to each DNA fragment size, from the whole sample of sequenced reads.

By default, *plot2DO* aligns the TSSs, and plots the coverage obtained by stacking the footprints of the of the entire fragments that were sequenced (Fig. S1). One may want to plot the distribution of the DNA fragments at other locations from the genome *{e.g*. +1 nucleosomes), or to plot the distribution of the fragment centers (nucleosome dyads) instead of the coverage of the entire fragments. When multiple samples are compared, the color scale needs to be specified in a consistent way for all plots. These options are all possible using the following command line flags:

~~~
$ Rscript plot2DO.R -file=yeast_SRR2045610.bam -type=dyadsreference=Plusl -colorScaleMax=0.15
$ Rscript plot2DO.R -f yeast_SRR2045610.bam -t dyads -r Plusl - 0.15          # short version
~~~

This command will generate Figure S2.

**Supplementary Figure S2.**
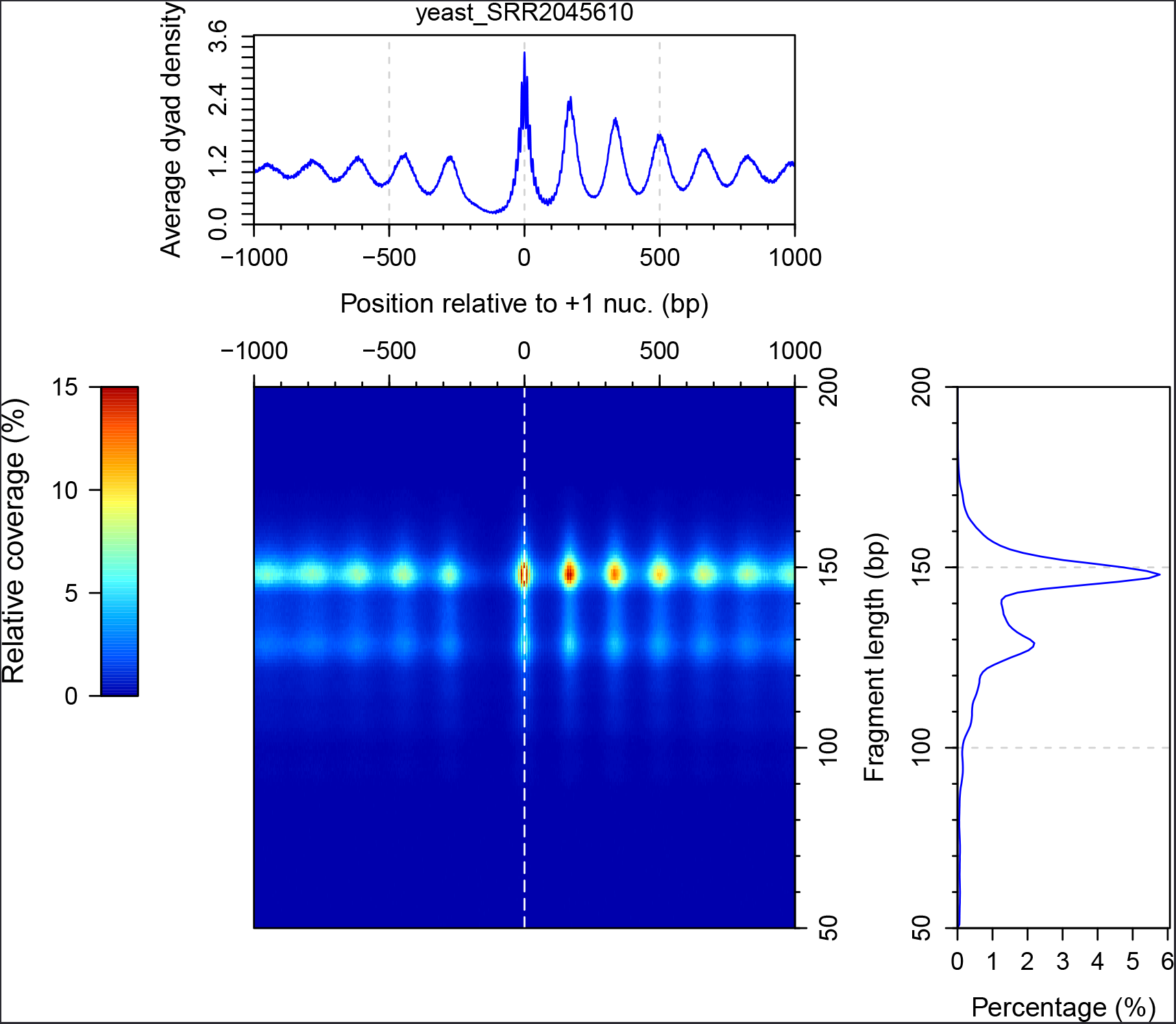
Distribution of nucleosome dyads near the typical positions of +1 nucleosomes. This special case of 2DO plot is also known as a V-plot (Henikoff et al., 2011).

Apart from the usual alignments, using the TSS, TTS or +1 nucleosomes as reference points, one can also align specific lists of reference points, provided in the form of a BED file. For convenience, we provide some of the yeast features that one may want to examine (tRNA genes, origins of replication, centromeres), and below are a few examples of how one can use *plot2DO* to inspect a specific list of loci.

~~~
$ Rscript plot2DO.R—f yeast_SRR2045610.bam —sites=Annotations/Yeast_tRNA_genes.bed —align=fivePrime
          — site Labe1=*t* RNA — simplifyPlot=on
$ Rscript plot2DO.R—f yeast_SRR2045610.bam —sites=Annotations/Yeast_ ARS_locations.bed —align=center
          — site Labe1=ARS — simp1ify P1ot=on
$ Rscript plot2DO.R—f yeast_SRR2045610.bam —sites=Annotations/ Yeast_centromeres.bed —align=threePrime
          — siteLabel=centromere —simplifyPlot=on
~~~

Notice that using the `-simplifyPlot=on` option, it is possible to plot only the 2DO panel (without the panels showing the one-dimensional occupancy and the histogram of fragment lengths), to make it easier to combine multiple such panels in a single figure (Fig. S3).

**Supplementary Figure S3.**
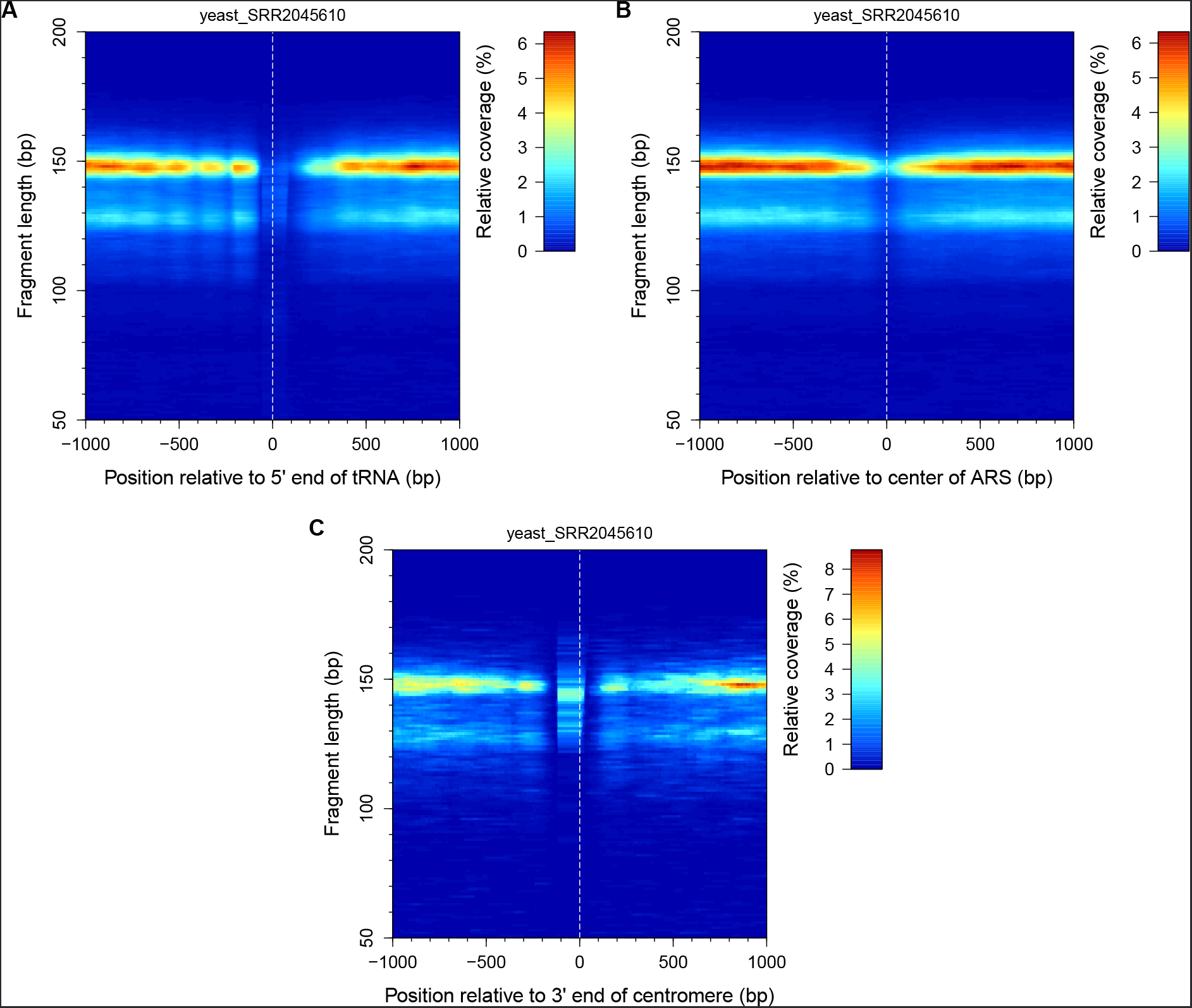
2DO plots showing the distribution of sequencing data near the yeast tRNA genes (A), origins of replication (B), and centromeres (C).

*Plot2DO* can process sequencing data from multiple organisms: *Saccharomyces cerevisiae, Drosophila melanogaster, Caenorhabditis elegans, Mus musculus* and *Homo sapiens*. The following genome versions are available for these organisms: yeast - sacCer3; fly - dm3, dm6; worm - celO, cell; mouse - mm9, mmlO; human - hgl8, hgl9. For the multicellular organisms the alignment of +1 nucleosomes is not possible, as the locations of these nucleosomes could vary from cell type to cell type, and these positions should be identified separately in each cell type. Here are a few examples of commands used to examine the distribution of reads at the TSSs of the aforementioned higher organisms.

~~~
$ Rscript plot2DO.R —file=fly_SRR2038265.bam —organism=dm3 —simplify Plot=on
$ Rscript plot2DO.R —file=worm_SRR3289717.bam —organism=ce10 —simplify Plot=on
$ Rscript plot2DO.R —file=mouse_SRR572708.bam —organism=mm10 —simplify Plot=on
$ Rscript plot2DO.R —file=human_SRR1781839.bam —organism=hg19 —simplify Plot=on
~~~

The figures resulted from these commands are shown in Figure S4.

**Supplementary Figure S4.**
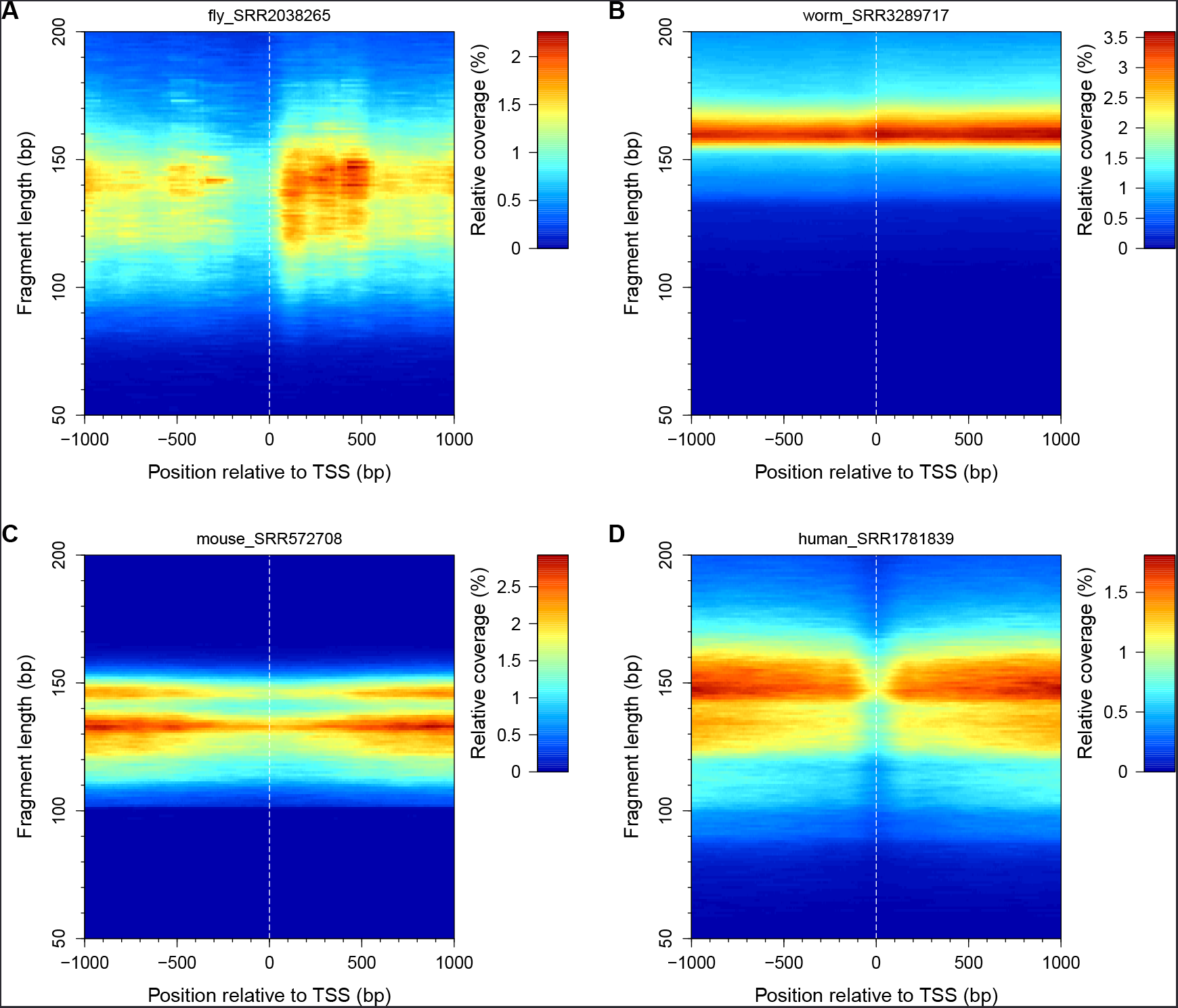
2DO plots showing the distribution of sequencing data near the TSS in fly (A), worm (B), mouse (C), and human (D).

If one wants to check only a small genomic region (*e.g*. all tRNA genes or +1 nucleosomes), the -squeezePlot=on option is very handy. Below are a few examples:

~~~
$ Rscript plot2DO.R -f yeast_50U_MNase_SRR3649301.bam -s Annotations/Yeast_tRNA_genes.bed —upstream =100
       —downstream=100 —squeezePlot=on
$ Rscript plot2DO.R -f yeast_100U_MNase_SRR3649296 .bam -s Annotations/Yeast_tRNA_genes.bed —upstream =100
       —downstream=100 —squeeze Plot=on
$ Rscript plot2DO.R -f yeast_200U_MNase_SRR3649297.bam -s Annotations/Yeast_tRNA_genes.bed —upstream =100
       —downstream =100 —squeeze Plot=on
$ Rscript plot2DO.R -f yeast_300U_MNase_SRR3649298.bam -s Annotations/Yeast_tRNA_genes.bed —upstream =100
       —downstream=100 —squeeze Plot=on
$ Rscript plot2DO.R -f yeast_400U_MNase_SRR3649299.bam -s Annotations/Yeast_tRNA_genes.bed —upstream =100
       —downstream=100 —squeeze Plot=on
$ Rscript plot2DO.R -f yeast_50U_MNase_SRR3649301.bam r Plusl -u 100 -d 100 —squeezePlot=on
$ Rscript plot2DO.R -f yeast_100U_MNase_SRR3649296.ba- r Plusl -u 100 -d 10 —squeezePlot=on
$ Rscript plot2DO.R -f yeast_200U_MNase_SRR3649297.bam -r Plus1 -u 100 -d 100 —squeezePlot=on
$ Rscript plot2DO.R -f yeast_300U_MNase_SRR3649298.bam -r Plus1 -u 100 -d 100 —squeezePlot=on
$ Rscript plot2DO.R -f yeast_400U_MNase_SRR3649299.bam -r Plus1 -u 100 -d 100 —squeezePlot=on
~~~

The figures resulted from these commands are shown as panels in Figure S5.

**Supplementary Figure S5.**
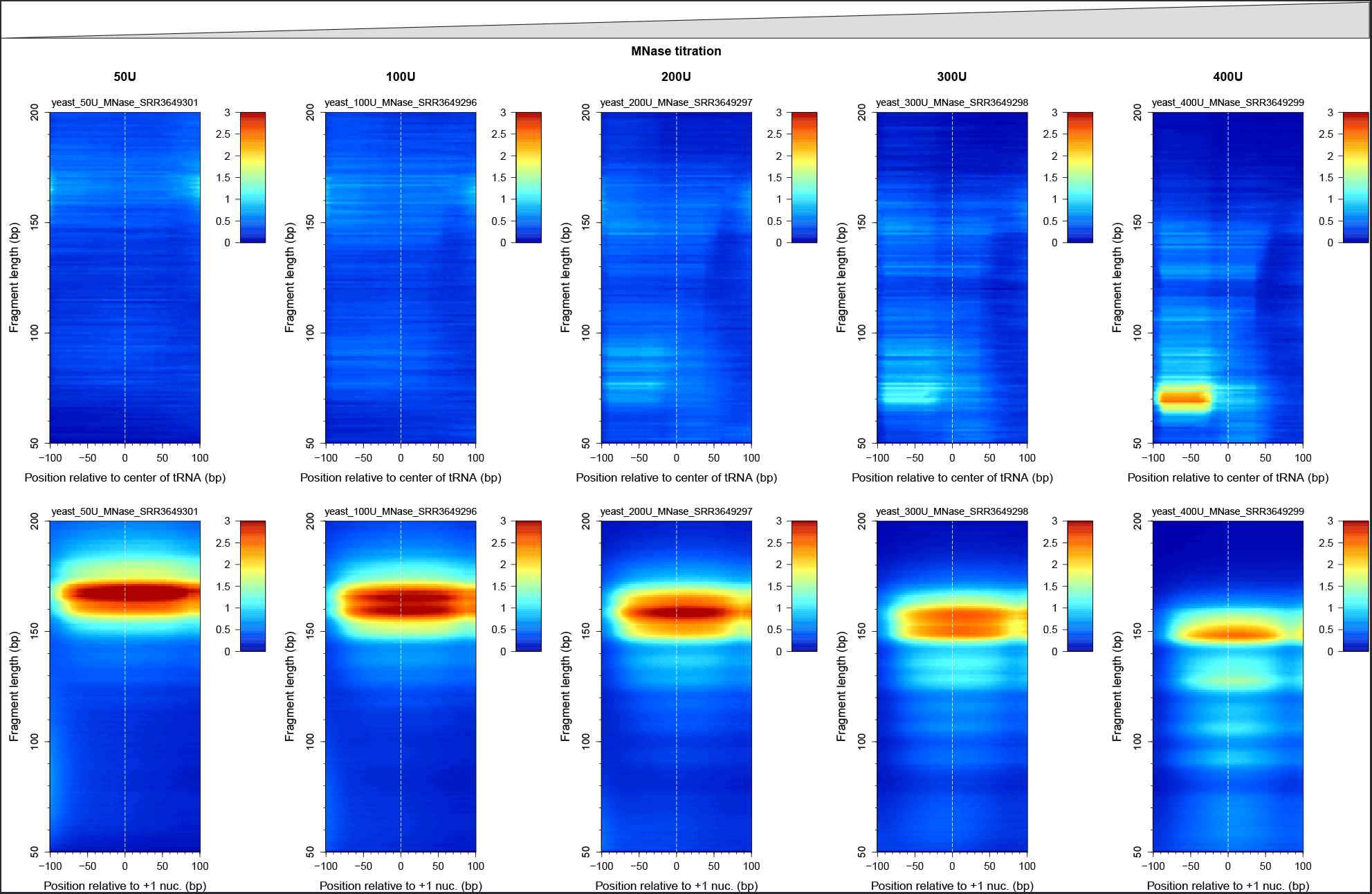
A titration of MNase produces different levels of chromatin digestion, and different sizes for the undigested fragments of DNA. Plus 1 nucleosomes are particularly sensitive to MNase, as they are located next to the nucleosome free region (the most accessible regions of the genome), while tRNA genes are more resistant to MNase.

To list all the available options of *plot2DO*, run the following command in the terminal:

~~~
$ Rscript plot2DO.R —help
Usage: plot2DO .R [options]

Options:
         -f FILE, -file=FILE
                Dataset file name [options: BED or BAM format]
         -t TYPE, -type=TYPE
                Types of distribution to plot [options: occ , dyads, fivePrime_ends , threePrime_ends;
   default = occ]
         -o ORGANISM,organism ORGANTSM
                Geno- version [options: sacCer3 (default), dm3, dm6, celO , cell, mm9, mmlO, hgl8 , hgl9]
         -r REFERENCE, -reference=REFERENCE
                Reference points to align [options: TSS (default), TTS, Plusl]
         -s SITES, -sites=SITES
                Reference points in BED format
         -a ALIGN, -align=ALIGN
                What points of the provided intervals to align? [options: center (default), fivePrime,
    threePrime]
        —siteLabe1=SITELABEL
                Label for the aligned sites [default = Sites]
         -1 MINLENGTH, -minLength=MINLENGTH
                The smallest DNA fragment to be considered [default = 50]
         -L MAXLENGTH, -maxLength VIAXLENGIH
                The largest DNA fragment to be considered [default = 200]
         -u UPSTREAM, -upstream=UPSTREAM
                Length of the upstream region that will be plotted [default = 1000]
         -d DOWNSTREAM, -downstreanrfOWNSTREAM
                Length of the downstream region that will be plotted [default = 1000]
         -m COLORSCALEMAX, -colorScaleMax=COLORSCALEMAX
                Maximum value on the color scale (e.g. 0.02)
        —simplifyPlot=SIMPLIFYPLOT
                Simplify the plot (show only the 2D Occ.) [options: on, off (default)]
         - squeezePlot=SQUEEZEPLOT
                Simplify the plot (show only the 2D Occ.) and squeeze the heat map [options: on, off (default)]
         -h , —help
                Show this help message and exit
~~~

### Data used in examples

The data that were used in the examples above were downloaded from GEO database. The corresponding accession numbers are listed below:

| Filename | GEO accession number for the raw data | Publication |
| --- | --- | --- |
| yeast_SRR2045610.bam | GSM1700669 | Ocampo et al. (2016) |
| yeast_50U_MNase_SRR3649301.bam | GSM2193286 | Chereji et al. (2017) |
| yeast_100U_MNase_SRR3649296.bam | GSM2193281 | Chereji et al. (2017) |
| yeast_200U_MNase_SRR3649297.bam | GSM2193282 | Chereji et al. (2017) |
| yeast_300U_MNase_SRR3649298.bam | GSM2193283 | Chereji et al. (2017) |
| yeast_400U_MNase_SRR3649299.bam | GSM2193284 | Chereji et al. (2017) |
| fly_SRR2038265.bam | GSM1694824 | Chereji et al. (2016) |
| worm_SRR3289717.bam | GSM2098437 | Jeffers and Lieb (2017) |
| mouse_SRR572708.bam | GSM1004654 | Teif et al. (2012) |
| human_SRR1781839.bam | GSM1425454 | West et al. (2014) |

